# The importance of UBQLN2 ubiquitylation for its turnover and localization

**DOI:** 10.1101/2025.10.02.679934

**Authors:** Martin Grønbæk-Thygesen, Caroline Kampmeyer, Paula Eschger, Michael H. Tatham, Marloes Arts, Kay Hofmann, Kresten Lindorff-Larsen, Wouter Boomsma, Rasmus Hartmann-Petersen

**Affiliations:** Department of Biology, University of Copenhagen, Ole Maaløes Vej 5, DK-2200 Copenhagen, Denmark; Institute for Genetics, University of Cologne, Cologne, Germany; Centre for Molecular, Cell and Developmental Biology, School of Life Sciences, University of Dundee, Dow Street, Dundee, DD1 5EH. UK; Department of Computer Science, University of Copenhagen, Copenhagen, Denmark

**Keywords:** IDR, IDP, LLPS, liquid-liquid phase separation, proteostasis, lysine desert

## Abstract

UBQLN2 is a member of the UBL-UBA domain protein family that functions as extrinsic substrate receptors for the 26S proteasome. UBQLN2 has been shown to undergo phase separation *in vitro*. In cells, UBQLN2 forms condensates that may be of importance for tuning protein degradation via the ubiquitin-proteasome system and potentially of relevance for *UBQLN2*-linked amyotrophic lateral sclerosis (ALS). Here we show that UBQLN2 is ubiquitylated on lysine residues in the N-terminal UBL domain. The C-terminal region of UBQLN2 is lysine-depleted, and we show that introducing lysine residues in this region leads to its E6AP-dependent degradation. The UBL domain critically stabilizes UBQLN2 and protects it from proteasomal degradation. Fusion of ubiquitin to the UBQLN2 N-terminus stabilizes UBQLN2 and increases its propensity for locating in puncta, indicating that ubiquitylation of the UBQLN2 UBL domain regulates abundance and localization.

## Introduction

Most intracellular proteins are degraded by the 26S proteasome, an abundant protease complex found in the cytosol and nucleus of all eukaryotic cells ^1-3^. Although the proteasome can degrade some proteins directly ^4^, most proteasome targets are first covalently conjugated to ubiquitin. This process, termed ubiquitylation ^1^, is catalyzed by an enzymatic cascade involving a ubiquitin-activating enzyme (E1), a ubiquitin-conjugating enzyme (E2) and a ubiquitin-protein ligase (E3). Typically, this results in the ε-amino group of a lysine residue in the target protein being conjugated to the C-terminal carboxy group in ubiquitin via an isopeptide bond. However, ubiquitylation of the N-terminal amino group via a peptide bond, serine and threonine residues via ester bonds, and cysteine residues via thioester bonds have also been observed ^5,6^. Additional cycles of ubiquitylation result in the target protein being conjugated to a poly-ubiquitin chain. As ubiquitin contains seven lysine residues, distinct types of ubiquitin chains can be formed. For proteasomal degradation, the principal signal is K48-linked ubiquitin chains, while other chain types have been linked to both proteasomal and non-proteasomal functions ^7,8^.

Ubiquitylation of the target protein enhances its affinity for intrinsic ubiquitin-binding subunits of the 26S proteasome ^9,10^. However, for efficient degradation, the 26S proteasome also requires so-called substrate shuttle proteins ^11-13^. These proteins, including RAD23 ^14^ and the ubiquilins (UBQLN1-4), contain N-terminal ubiquitin-like (UBL) domains that interact with the 26S proteasome ^15-18^ and C-terminal ubiquitin-associated (UBA) domains that bind ubiquitin chains ^17,18^. This constellation of domains thus allows the substrate shuttles to function as extrinsic proteasomal ubiquitin receptors ^11-13,18^. However, importantly, the UBQLNs have also been shown to mediate ubiquitin-independent proteasomal degradation of certain proteins ^4^. Presumably, due to their interactions with the E3s ^19,20^ and the 26S proteasome ^15,18,21-23^, the UBL-UBA proteins have evolved long stretches that are depleted for lysine residues ^24,25^, thus protecting them from untimely ubiquitylation and degradation ^24^. In UBQLN2, this so-called lysine desert covers more than 500 residues and is found in orthologues from yeast to man.

Intriguingly, UBQLN2 has been shown to phase separate *in vitro* at physiological concentrations and form condensates in cells ^26-28^, and can be incorporated into stress granules ^29^. The endogenous phase separation properties of UBQLN2 are dependent on its STI1-3/4 regions and the UBA domain located in the lysine desert of UBQLN2, and is also regulated by ubiquitin-binding ^26,30^ and proposed to be promoted by E3s ^31^. Thus, polyubiquitylated proteins promote phase separation, and are incorporated into condensates, which may recruit proteasomes ^32-35^ to control protein degradation ^36^, and *in vitro* UBQLN2 also recruits the E3 ligase E6AP into condensates ^20^.

Notably, gene variants in *UBQLN2* have been linked to a dominant X-linked form of amyotrophic lateral sclerosis (ALS) ^37-42^. Several of the pathogenic variants cluster in a proline-rich stretch within the disordered lysine-desert region, and various mechanisms for how these variants affect UBQLN2 function have been proposed ^40,42-44^.

Although the role of non-covalent ubiquitin-binding to the UBQLN2 UBA domain on phase separation has been studied in detail ^26,30^, not much is known about the effect of covalent ubiquitin binding (ubiquitylation). Here we describe cellular studies on the ubiquitylation of UBQLN2. We show that UBQLN2 is ubiquitylated on lysine residues in the N-terminal UBL domain. The UBL domain critically stabilizes UBQLN2 and protects it from ubiquitin-independent proteasomal degradation. Introducing lysine residues in the UBQLN2 lysine desert leads to E6AP-catalyzed UBQLN2 ubiquitylation and proteasomal degradation. Finally, we show that artificial fusion of ubiquitin to the UBQLN2 N-terminus increases its abundance and promotes its localization in cytosolic puncta.

## Results

### UBQLN2 is a lysine and cysteine depleted protein

Previously, we performed a bioinformatics screening for proteins containing long stretches devoid of lysine residues ^24,45^. This led to the identification of hundreds of so-called lysine-desert proteins, including UBQLN1, UBQLN2 and UBQLN4. Although UBQLN3 is also strongly lysine depleted, it does contain three lysine residues downstream of the UBL domain that are not found in the other UBQLN paralogs (**Fig. S1**). None of the UBQLNs show similar underrepresentation of arginine (**Fig. S1**), suggesting a specific selection against lysine rather than evolutionary pressure to avert positive charges. Moreover, in addition to lysine, the UBQLNs are also highly depleted for cysteine (the cysteine frequency in the human reference proteome is 2.3%), with UBQLN1 and UBQLN2 not containing any cysteine residues at all (**Fig. S1**). UBQLN2 has been extensively studied and gene variants have been linked to amyotrophic lateral sclerosis and frontotemporal dementia (ALS/FTD) ^37-41^, and thus we decided to explore the role of the lysine desert in this particular paralogue.

Multiple sequence alignments revealed that the positioning of the lysine residues in UBQLN2 orthologues is conserved and restricted to the UBL domain (**Fig. 1A**). To further evaluate if evolution has selected against introducing particular amino acids into specific regions of UBQLN2, we applied the ESM-2 protein language model ^46^ which has previously been shown to be sensitive to the impact of individual amino acid substitutions on protein structure and function ^46^. In agreement with our manual sequence alignments (**Fig. 1A, Fig. S1**), lysine (and cysteine) residues outside the UBL domain were strongly disfavored by the ESM-2 model (**Fig. 1B, SupplementalFile**), suggesting evolutionary pressure disfavoring these residues in UBQLN2. According to structural models, except for the conserved K58 (relative accessible surface area, rASA = 0.07) and K38 (rASA = 0.39), the other lysine residues of the UBL domain are exposed (rASA ≥ 0.5) (**Fig. 1C**).

**Fig. 1.**
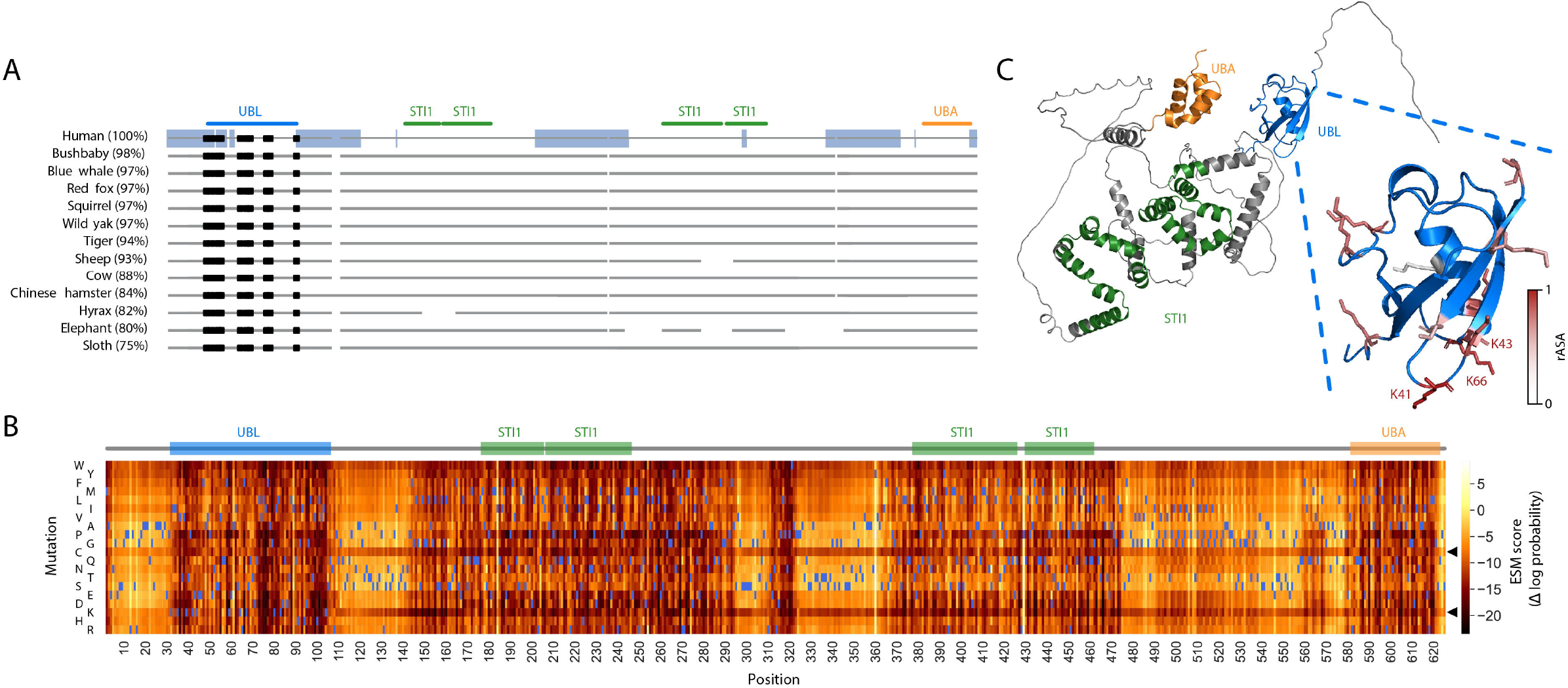
UBQLN2 is a conserved lysine-depleted protein. (A) Sequence comparison of UBQLN2 orthologs in the indicated species. Intrinsically disordered regions in human UBQLN2 based on MobiDB are shown as a blue bar. The domain organization based on the SMART database is marked. Lysine residues are marked as black squares. (B) ESM-2 predictions of all possible single amino acid substitutions of human UBQLN2 presented as a heat map. The wild-type residues are marked in blue. ESM-2 scores close to zero (light yellow colors) indicate that the amino acid substitution is compatible with the ESM-2 language model, whereas negative scores (dark orange colors) indicate that the variant is incompatible with the ESM-2 model. The domain organization (based on SMART) is aligned above the map. Note that substitutions to lysine or cysteine in general appear detrimental, in particular downstream of the UBL domain. (C) The AlphaFold2 predicted structured of human UBQLN2 (AF-Q9UHD9-F1) (left panel). The UBL domain is colored blue, and the UBA domain is colored orange and the STI1 regions green. Zoom in on the UBL domain (right panel) with the lysine residues highlighted as stick representations and colored based on the relative accessible surface area (rASA, dark red exposed; grey, buried).

### UBQLN2 is ubiquitylated on lysine residues in the UBL domain

To test if UBQLN2 is subject to ubiquitylation, we first expressed wild-type UBQLN2 carrying a C-terminal HA-tag together with N-terminally myc-tagged ubiquitin in HEK293T cells. The cells were treated with the proteasome inhibitor bortezomib to block degradation of ubiquitylated proteins. To purify proteins covalently conjugated to ubiquitin, the cell lysates were first denatured by incubation at 100 °C in SDS. Then, the SDS was neutralized with Triton X-100 and the tagged ubiquitin was immunoprecipitated. Western blotting of the precipitated material revealed the presence of ubiquitylated UBQLN2 (**Fig. 2A**).

**Fig. 2.**
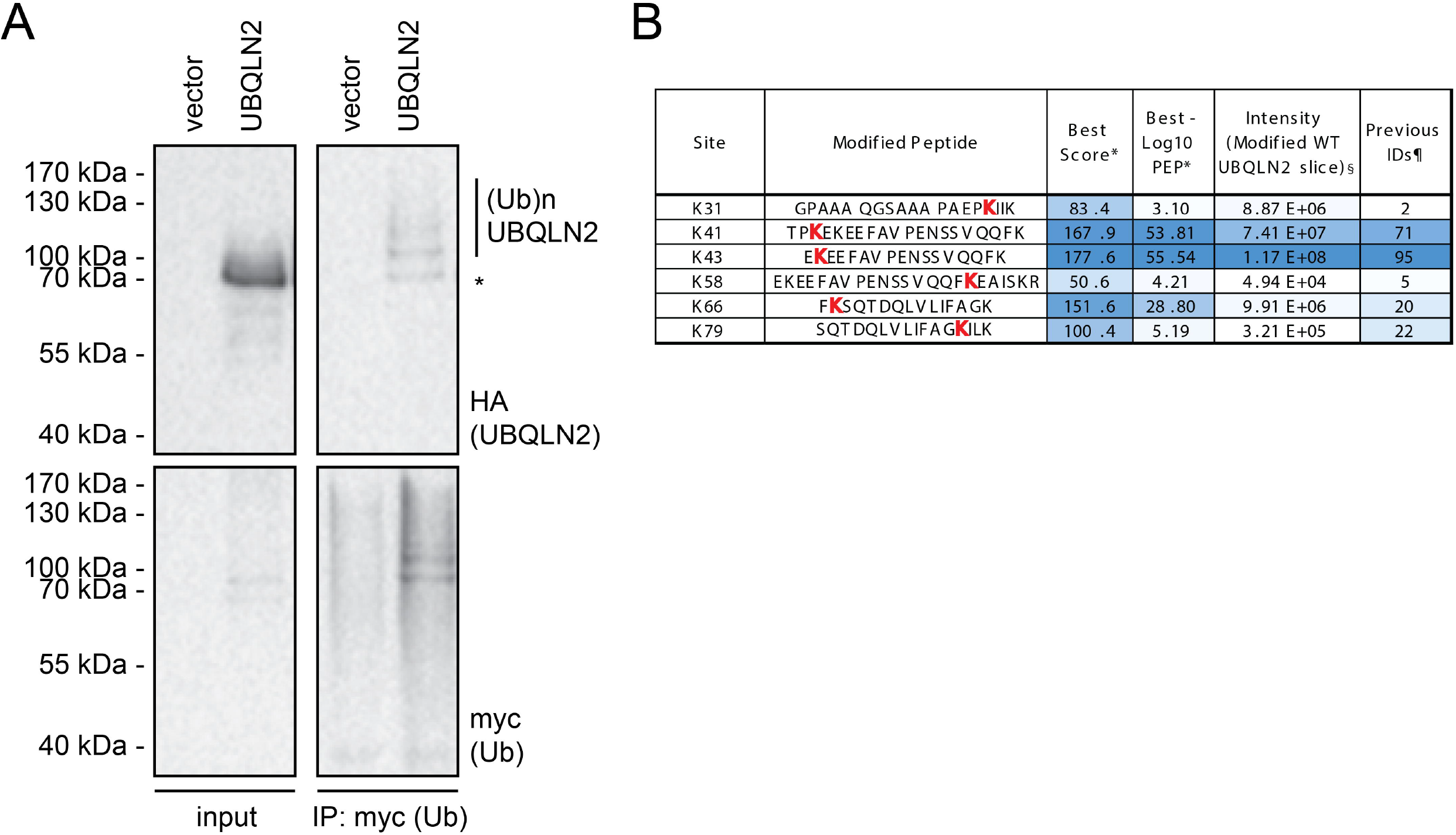
UBQLN2 is ubiquitylated on UBL-domain lysine residues. (A) HEK293T cells transiently co-transfected with plasmids expressing myc-ubiquitin and either vector (control) or wild-type UBQLN2-HA were treated with 10 µM bortezomib (BZ) for 16 hours. Following denaturing immunoprecipitation (IP) of myc-tagged ubiquitin, the proteins were analyzed by SDS-PAGE and western blotting. Ubiquitylated proteins are marked. The asterisk (*) marks the unmodified protein. (B) Summary of sites of ubiquitylation identified in HA-tagged UBQLN2 immunoprecipitated from HEK293T cells. *, Best data for either GG-K or LRGG-K modified peptides for the same site (**Fig. S4, Fig. S5**). §, Summed intensity of both GG-K and LRGG-K peptides for the same site derived from the modified WT UBQLN2 slice (**Fig. S2A**). ¶, Number of times site is reported on Phosphosite.org (accessed 05/01/2025).

To identify the sites in UBQLN2 that are ubiquitylated, cell lysates were subjected to denaturing immunoprecipitation of the HA-tagged UBQLN2 and analyzed by SDS-PAGE and mass spectrometry (**Fig. S2**). Wild-type UBQLN2 contains 11 lysines, for which 6 were reported by MaxQuant with reasonable evidence of modification by either GG-K or LRGG-K (one missed cleavage) adducts (**Fig. 2B**). The strongest evidence for ubiquitylation was found for lysines 41, 43, and 66, which cluster together on one face of the UBL domain predicted to orient towards the remainder of the protein (**Fig. 1C**). There was weaker evidence for modification at K79, K31 and K58 (**Fig. 2B**). These results are consistent with high thought-put proteomics studies reported on PhosphoSitePlus ^47^ for which the bulk of the evidence identifies lysines 41 and 43 as major acceptors, with lysines 66 and 79 being identified less frequently (**Fig. 2B**). Evidence for modification of the remaining lysines is rare (https://www.phosphosite.org/proteinAction.action?id=737 - accessed 03/01/2025). The MS/MS spectra (**Fig. S3, Fig. S4**) and data (**SupplementalFile**) are included in the supplemental material.

### E6AP and proteasome dependent degradation of a lysine-rich UBQLN2 variant

As a next step in our characterization of UBQLN2 ubiquitylation, we performed experiments similar to the ones we previously carried out for other lysine desert proteins ^24^. Wild-type UBQLN2, as well as a UBQLN2 version, where multiple arginine residues had been substituted for lysine (R□→□K) (**Fig. S5**), were introduced into HEK293T cells along with myc-tagged ubiquitin. As a control, we also included a UBQLN2 variant, where the arginine residues were replaced with glutamine (R□→□Q) (**Fig. S5**). Denaturing immunoprecipitations from cultures treated with bortezomib (as above) confirmed ubiquitylation of wild-type UBQLN2 (**Fig. 3A**). However, the ubiquitylation was more pronounced for the R→K mutant **(Fig. 3A**) and reduced in the glutamine control **(Fig. 3A**). The reduced modification of the R□→□Q variant was unexpected but indicates that the arginines in the disordered region affects ubiquitylation of the UBL domain lysine residues, possibly via perturbing the localization of UBQLN2 or its ability to form puncta (see below). In addition, we noted that the unmodified R□→□Q variant migrated slightly slower on the gel compared to wild-type and R□→□K.

**Fig. 3.**
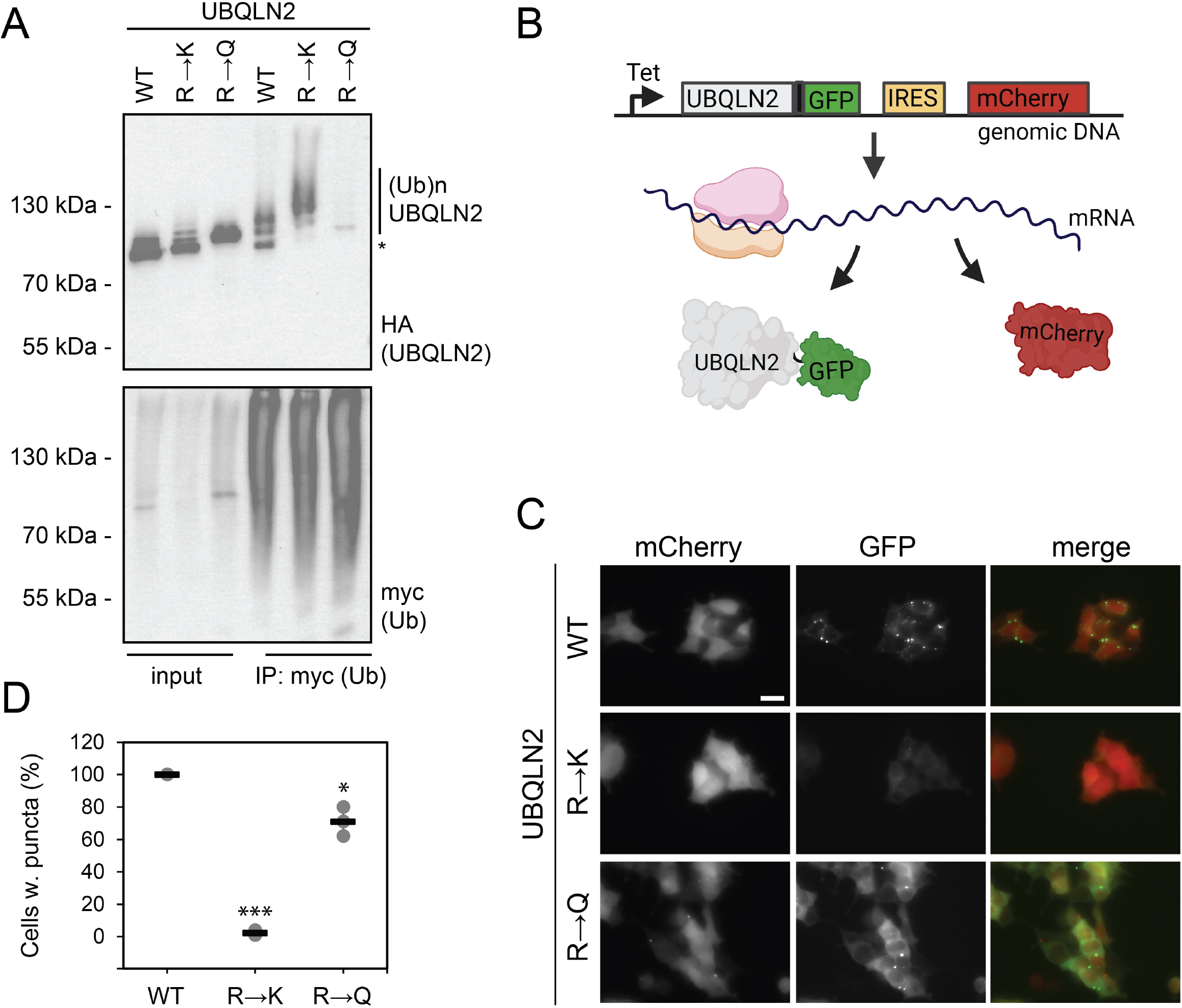
Introduction of lysine residues leads to increased UBQLN2 ubiquitylation. (A) HEK293T cells transiently co-transfected with expression constructs for myc-tagged ubiquitin and UBQLN2 (C-terminally HA-tagged) WT, RL→LK and RL→LQ, were treated with 10 µM bortezomib (BZ) for 16 hours. Following denaturing immunoprecipitation (IP) of myc-tagged ubiquitin, the proteins were analyzed by SDS-PAGE and western blotting. Ubiquitylated proteins are marked. The asterisk (*) marks the unmodified protein. (B) Schematic illustration of the expression system used for producing UBQLN2-GFP variants. Note that mCherry is produced from an internal ribosomal entry site (IRES) in the same mRNA as UBQLN2-GFP. Created in BioRender. Hartmann-Petersen, R. (2025) https://BioRender.com/0cya3n6. (C) Representative fluorescence micrographs of live HEK293T cells expressing the indicated constructs. The scale bar represents 20 µm. (D) Cells expressing the indicated UBQLN2-GFP fusions containing puncta were manually quantified by fluorescence microscopy. The frequency of cells containing puncta in independent experiments are represented as data points (minimum of 100 mCherry positive cells, n = 3). MS Excel was used to analyze data using a t-test (*, p= 0.031; ***, p<0.001). Means (horizontal line) and standard deviations (error bars) are shown.

To assess how the introduction of lysine affected the subcellular localization and abundance of UBQLN2, we generated stable transfectants expressing the UBQLN2 variants fused to GFP from a single genomic locus. To control for cell-to-cell variations in expression, mCherry was produced from an internal ribosomal entry site (IRES) in the same mRNA as UBQLN2-GFP (**Fig. 3B**). Live-cell fluorescence microscopy revealed a punctate-like localization of wild-type UBQLN2-GFP (**Fig. 3C**) in all cells (**Fig. 3D**), consistent with previously observed condensates ^26-28^. In comparison, the level of the R□→□K variant and puncta formation was strongly reduced. The R□→□Q variant expression appeared to be higher than WT (**Fig. 3C**) and did form puncta (**Fig. 3D**). In addition, the R□→□Q variant displayed a wider cytosolic distribution (**Fig. 3C**), indicating that the arginine residues directly or indirectly also contribute in tuning UBQLN2 abundance and localization. The effect of the mutations on UBQLN2 abundance were confirmed by western blotting (**Fig. 4A**). Treatment of cells with the proteasome inhibitor bortezomib (BZ) increased the abundance of WT UBQLN2 modestly, and the R→K mutant strongly, while the R→Q variant appeared not to be affected by proteasome inhibition (**Fig. 4A**). This suggests that ubiquitylation of lysine residues in the lysine desert leads to proteasomal degradation of the UBQLN2 R□→□K variant.

**Fig. 4.**
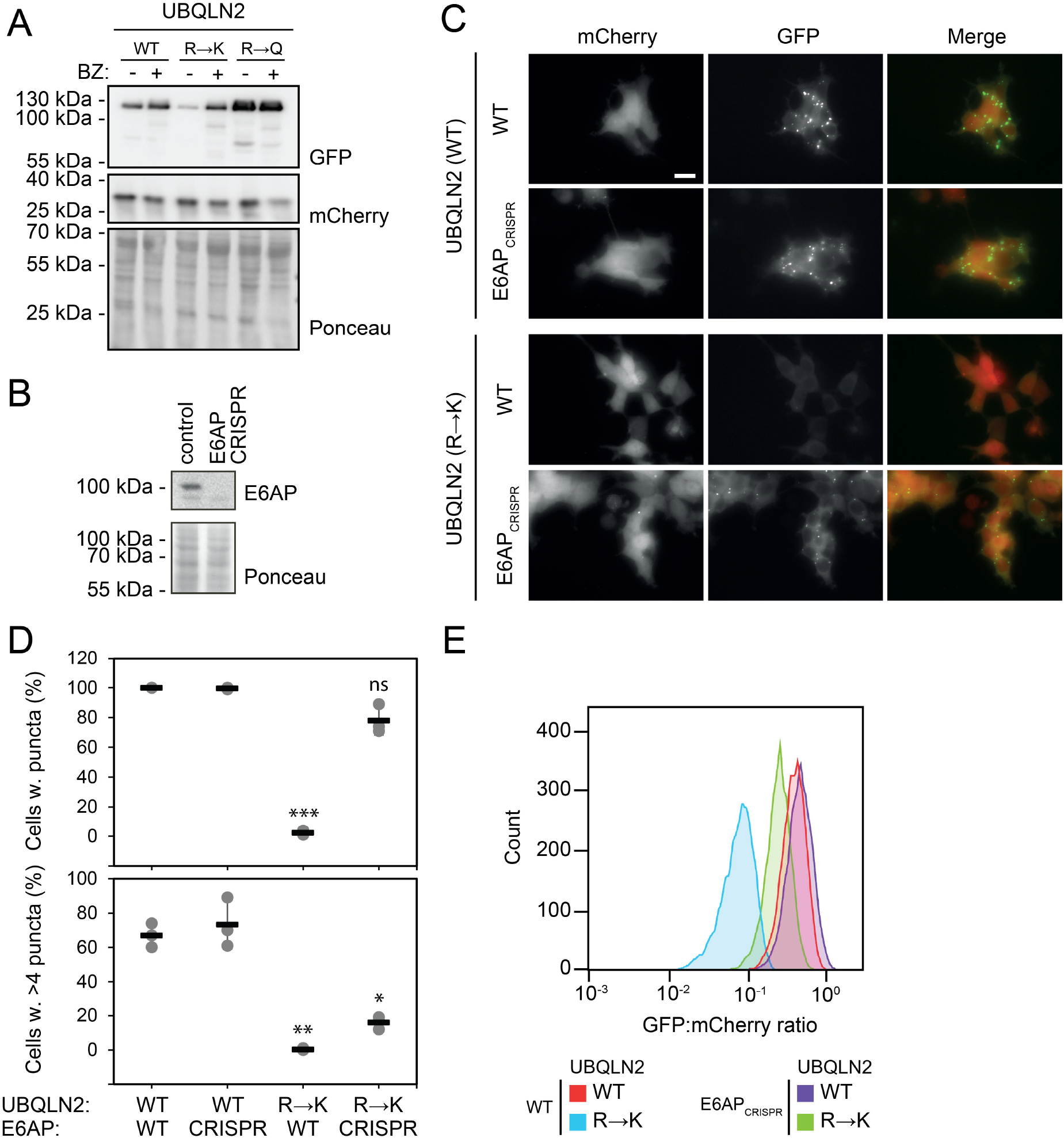
E6AP is required for ubiquitylation of the UBQLN2 lysine variant. (A) HEK293T cells stably transfected to express the indicated UBQLN2-GFP variants were treated (+) or untreated (-) with the proteasome inhibitor bortezomib (BZ) for 8 hours and analyzed by SDS-PAGE and western blotting. A Ponceau S staining and blotting for mCherry were included as controls. (B) Whole cell lysates of HEK293T cells, wild-type (control) or CRISPR disrupted for E6AP, were analyzed by SDS-PAGE and western blotted for E6AP. A Ponceau S staining was included as a control for equal loading. (C) Representative fluorescence micrographs of live HEK293T landing pad cells, either wild-type (WT) or CRISPR disrupted for E6AP, expressing the indicated UBQLN2-GFP variants. The scale bar represents 20 µm. (D) Wild-type (WT) and E6AP CRISPR cells expressing the indicated UBQLN2-GFP fusions containing puncta were manually quantified by fluorescence microscopy. The frequency of cells containing puncta in independent experiments are represented as data points (minimum of 100 mCherry positive cells, n = 3). MS Excel was used to analyze data using a t-test (ns, not significant; *, p=0.139; **, p=0.003;***, p<0.001). Means (horizontal line) and standard deviations (error bars) are shown. (E) Flow cytometry-based quantification of the GFP-tagged proteins presented as histograms of the number of cells and the GFP:mCherry ratios. The mean and standard deviation (± SD) was for WT UBQLN2 in WT cells: 0.42±0.15, WT UBQLN2 in E6AP CRISPR cells: 0.50±0.20, RL→LK UBQLN2 in WT cells: 0.09±0.05, and RL→LK UBQLN2 in E6AP CRISPR cells: 0.27±0.11. A minimum of 35,000 cells were analyzed.

Since UBQLN2 has been shown to interact directly with the E3 ubiquitin-protein ligase E6AP ^20^, we decided to test if E6AP was responsible for the ubiquitylation and degradation of UBQLN2. To this end, we first generated HEK293T cells lacking E6AP using CRISPR (**Fig. 4B**) and then analyzed the subcellular localization of UBQLN2 by fluorescence microscopy. The localization of wild-type UBQLN2 was unaffected in the E6AP null cells, whereas for the R□→□K variant, levels were restored. In addition, puncta re-emerged in essentially all cells (**Fig. 4CD**), albeit they appeared smaller (**Fig. 4C**), and we observed fewer puncta per cell (**Fig. 4D**). Quantifying UBQLN2 abundance by flow cytometry also revealed that the abundance of the R□→□K variant was higher in the E6AP null cells, while the level of wild-type UBQLN2 was only very modestly increased with loss of E6AP (**Fig. 4E**).

### Fusion of ubiquitin to the UBL domain affects UBQLN2 abundance and localization

Next, we turned our attention to the UBL domain. Since we found that wild-type UBQLN2 was ubiquitylated on UBL domain lysine residues, we generated a UBQLN2 variant (K0) where all lysine residues were exchanged with arginine. This UBQLN2_K0_ mutant cannot form the canonical isopeptide bond linkages with ubiquitin. Conversely, to mimic ubiquitylated UBQLN2, we fused ubiquitin to the UBQLN2 N-terminus. We fused both wild-type ubiquitin and a lysine-less (K0) ubiquitin to wild-type and K0 UBQLN2. To avoid these fusions being cleaved by deubiquitylating enzymes, the two C-terminal glycine residues in ubiquitin were omitted (ΔGG). Finally, we generated a UBQLN2 variant lacking the UBL domain entirely (ΔUBL) and instead starting at position 108 (**Fig. 5A**). Transient expression of these C-terminally HA-tagged constructs followed by western blotting revealed the expected ubiquitylation pattern of the proteins (**Fig. 5B**), with a strong ubiquitin signal only being observed for UBQLN2 forms containing lysines. This was confirmed by mass spectrometry of the purified UBQLN2 proteins, which showed that only WT UBQLN2 constructs were strongly modified by ubiquitin (**Fig. S2A-D**). Notably, six lysines in ubiquitin were shown to be involved in ubiquitin-ubiquitin linkages (**Fig S2D**), although peptides indicative of these were predominantly detected in preparations containing the N-terminal ubiquitin fusion to UBQLN2, suggesting it is the fused ubiquitin which is modified. No evidence of N-terminal ubiquitylation of any UBQLN2 construct was detected in the mass spectrometry data. Thus, while lysine residues in both ubiquitin and the UBL domain are ubiquitylated, the lysine-less proteins (K0 and ΔUBL) were not measurably ubiquitylated, indicating that UBQLN2 is not susceptible to N-terminal ubiquitylation.

**Fig. 5.**
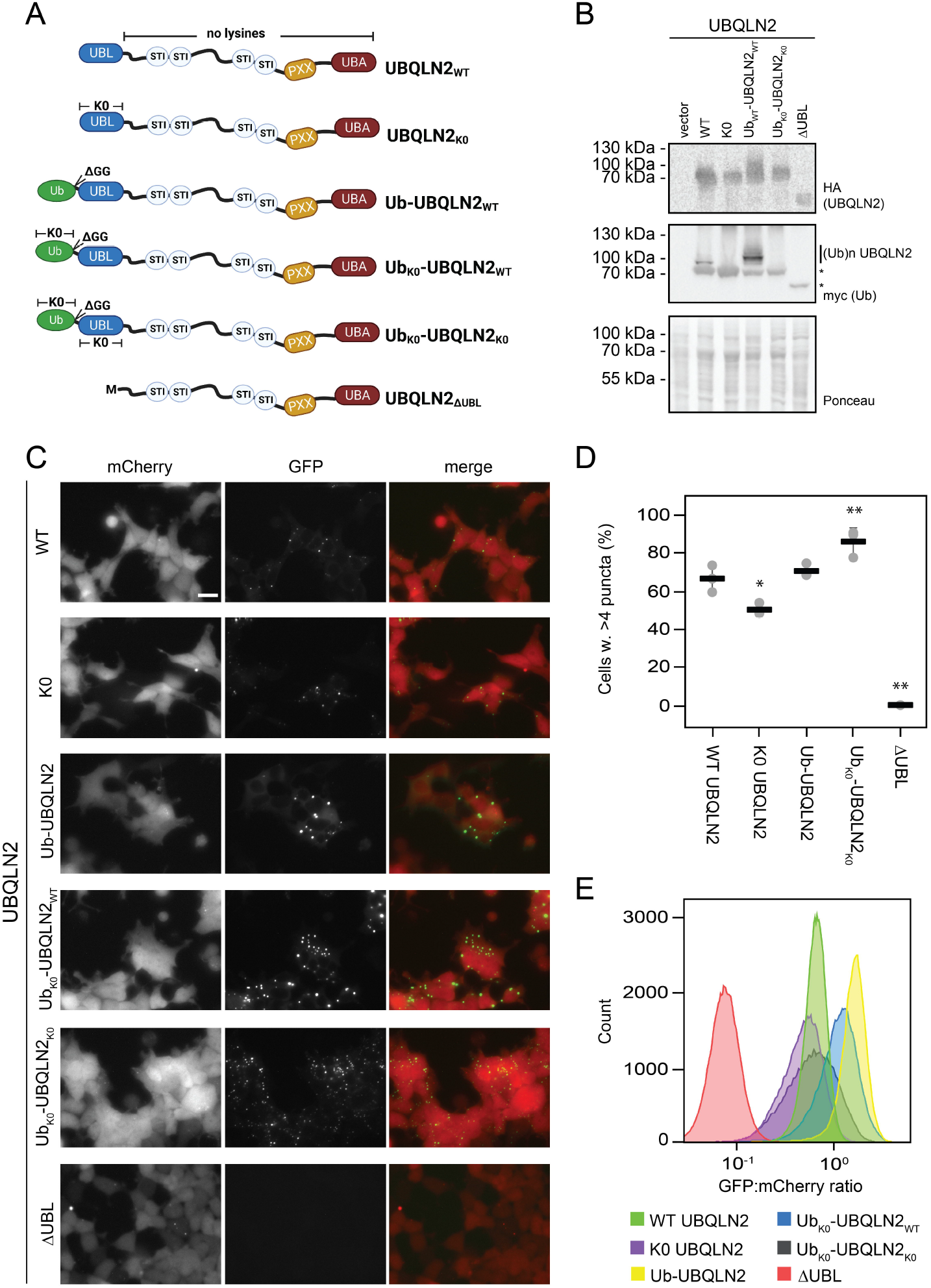
Artificial ubiquitin-fusion to UBQLN2 modulates the propensity for punctate formation. (A) Schematic illustration of the UBQLN2 variants used for the following experiments. Created in BioRender. Hartmann-Petersen, R. (2025) https://BioRender.com/qa447rp. (B) Whole cell lysates of HEK293T cells transiently transfected to express myc-ubiquitin and the indicated UBQLN2-HA variants were analyzed by SDS-PAGE and western blotting. Ponceau S staining serves as a control for equal loading. Ubiquitylated proteins are marked. The asterisks (*) mark the unmodified protein. (C) Representative fluorescence micrographs of live HEK293T cells expressing the indicated UBQLN2-GFP constructs. The scale bar represents 20 µm. (D) Cells expressing the indicated UBQLN2-GFP fusions containing >4 puncta per cell were manually quantified using fluorescence microscopy. The frequency of cells containing puncta in independent experiments are represented as data points (minimum of 100 mCherry positive cells, n = 3). MS Excel was used to analyze data using a t-test (*, p= 0.026; **, p=0.009 (UbK0-UBQLN2K0); **, p=0.004 (ΔUBL)). Means (horizontal line) and standard deviations (error bars) are shown. (E) Flow cytometry-based quantification of the GFP-tagged proteins presented as histograms of the number of cells and the GFP:mCherry ratios. The mean and standard deviation (±SD) was for WT UBQLN2: 0.23±0.11, K0 UBQLN2: 0.17±0.10, Ub-UBQLN2: 0.47±0.38, Ub(K0)-UBQLN2: 0.41±0.21, Ub(K0)-UBQLN2(K0): 0.26 ±0.16, and UBQLN2 ΔUBL: 0.08±0.04. A minimum of 58,000 cells were analyzed.

Stable transfectants, expressing the UBQLN2 variants fused to GFP, revealed that puncta formation was slightly reduced in the K0 variant, while it was more pronounced in the ubiquitin-UBQLN2 fusions (**Fig. 5CD**), potentially due to inter-protein ubiquitin-UBA interactions. Flow cytometry revealed some differences in the abundance of these UBQLN2 variants (**Fig. 5E**). For the Ub-UBQLN2 and Ub(K0)-UBQLN2 fusions the level appeared increased, which in case of Ub(K0)-UBQLN2 could explain the increased puncta formation. For the ΔUBL variant, the level was strongly reduced (**Fig. 5E**), which likely, at least in part, accounts for its strongly reduced puncta formation (**Fig. 5CD**).

Since the ΔUBL variant does not contain any lysine residues and was not observed to be ubiquitylated (**Fig. S2**), the reduced abundance is unlikely to depend on ubiquitylation. Accordingly, the level of the ΔUBL was unaffected by inhibiting the ubiquitin E1 with MLN7243 (**Fig. S6**). However, as the level of UBQLN2 ΔUBL clearly increased in response to proteasome inhibition (**Fig. S6**), we conclude that the ΔUBL protein is likely subject to ubiquitin-independent proteasomal degradation.

## Discussion

The ubiquilins, and UBQLN2 in particular, are intriguing and evolutionarily conserved proteins that are broadly involved in protein degradation. Through their N-terminal UBL domains, these proteins interact with the 26S proteasome, while their C-terminal UBA domains interact with ubiquitylated proteasome substrates ^16-18^. Between these folded domains, the proteins are predicted to be intrinsically disordered but do contain so-called STI1 regions involved in multimerization and condensate formation ^26,48^. In yeast cells, the single UBQLN ortholog, Dsk2, is critical for recruiting 26S proteasomes to cellular condensates during stress conditions ^32^. *In vitro*, human UBQLN1, UBQLN2 and UBQLN4 have been shown to undergo phase separation ^49^ and UBQLN2 condensates can recruit E6AP^20^.

Intriguingly, UBQLN1, UBQLN2, UBQLN4, and to a lesser extent also UBQLN3, contain long stretches devoid of both lysine and cysteine residues. As we show here for UBQLN2, this so-called lysine desert is critical for UBQLN2 to escape E6AP-catalyzed ubiquitylation and proteasomal degradation. Presumably, the lack of cysteine residues protects UBQLN2 from cysteine ubiquitylation and degradation, as recently shown for yeast Hrd1, another lysine- and cysteine-depleted disordered protein ^5^. However, since cysteine is also an unfavorable residue for disorder ^50^, this may also play a role.

We show that wild-type UBQLN2, similar to its ortholog in budding yeast, Dsk2 ^51^, is ubiquitylated on lysine residues in the UBL domain. These modifications are likely to affect proteasome-binding, turnover and puncta formation. However, since only a fraction of UBQLN2 is modified, it is not straightforward to test this directly. Still in support of this, we did observe a slight reduction in puncta formation of the UBQLN2 K0 variant and that artificial fusion of ubiquitin to the UBQLN2 N-terminus increases its localization in puncta. Thus, possibly ubiquitylation of the UBL domain increases UBQLN2 condensate formation. However, as abundance is also affected, the changes in puncta formation may be a consequence of the UBQLN2 levels. Indeed, UBQLN2 condensation has been noted to be sensitive to UBQLN2 levels ^26,37,52^. Moreover, N-terminal fusion of ubiquitin to UBQLN2 via a peptide bond is obviously topologically different than ligation to an ε-amino group of lysine side chain via an isopeptide bond, and we can therefore not rule-out that the increased puncta formation of the ubiquitin-UBQLN2 fusions is a consequence of the linear ubiquitin. Notably, in our mass spectrometry analysis we did not observe ubiquitylation of the N-terminal amino group in UBQLN2. For the ΔUBL truncation, puncta formation was strongly reduced. This could potentially be linked to the disordered N-terminal region upstream of the UBL domain, which is important for phase separation ^49^, but since the ΔUBL abundance is reduced it is possible that puncta formation would be apparent at higher expression levels. Therefore, it appears that separating puncta formation and abundance is not straightforward, and we only clearly achieve this for the R□→□Q variant.

Our analyses of the R□→□K variant in both wild-type and E6AP null cells suggest a complex effect on condensate formation. Arginine is known to be a stronger driver of phase separation than lysine ^53-56^, and thus R□→□K is expected to decrease the intrinsic propensity to form condensates. At the same time, the lysine variant shows increased ubiquitylation and lowered abundance in wild-type cells, which will also likely contribute to lowered puncta formation. The comparison between puncta formation of the lysine variant in wild-type and E6AP null cells suggests that the decreased formation of condensates in wild-type cells results from a combination of a direct effect of arginine on condensates and an effect of lowered abundance of the lysine variant.

N-terminal fusion of non-cleavable ubiquitin to proteins has been shown to result in rapid proteasomal degradation via the so-called ubiquitin-fusion degradation (UFD) pathway ^57-59^. This was not the case for our ubiquitin-UBQLN2 fusions, which may be connected with the stability of the C-terminal UBA domain. In the case of another UBL-UBA protein, RAD23, it has been shown that the cellular stability of this protein critically depends on the C-terminal UBA preventing initiation of its proteasomal degradation ^60-63^. We found that UBQLN2 lacking the UBL domain was rapidly degraded via the proteasome. Since this protein does not contain any lysine (or cysteine) residues, its degradation is likely independent of ubiquitin. However, due to the lack of the UBL domain, the protein should be unable to interact directly with the 26S proteasome, and we therefore speculate that it must interact with the proteasome via another mechanism, e.g. endogenous full-length UBQLN2 interacting via the STI1-3/4 regions and/or via ubiquitylated substrates. Since UBQLN2 is also connected to degradation via autophagy ^42,64-66^, this may also contribute to regulating UBQLN2 abundance, localization and function.

Finally, we note that the UBQLN2 variants in this study are all overexpressed fusion proteins. Since the UBQLN2 lysine residues are positioned in the N-terminal region and epitope tags placed on the N-terminus of the UBQLN2 can tune phase separation ^49^, we tagged the C-terminus of UBQLN2. However, this places the fusion partners in proximity to the small C-terminal UBA domain which could influence its activity. In addition, to enrich for ubiquitylated proteins our denaturing purifications were performed using cells treated with proteasome inhibitor, which will obviously affect overall cellular proteostasis.

## Materials and methods

### Bioinformatics

Homologue sequences were retrieved from UniProt and aligned using ClustalW (v.2.1). Disordered regions were extracted from MobiDB ^67^, and domains annotated according to the SMART database ^68^. For the heatmap, ESM-2 ^46^ was used to predict the log probability differences between the wildtype and all possible mutants per residue position. The structure visualizations were created using PyMOL, which was also used to calculate relative surface accessibility (rASA) scores (using the get_sasa_relative function).

### Plasmids

All plasmids were based on pcDNA3.1 (for transient transfections) or pVAMP ^69^ (for stable transfections). Cloning and mutagenesis was performed by Genscript. The plasmid for expression of myc-tagged ubiquitin has been described before ^24^.

### Cell culture

The HEK293T cells modified to contain a landing pad for site-specific integration ^69^ were cultured in Dulbecco’s modified Eagles medium (DMEM), containing 2 μg/mL doxycycline (Sigma-Aldrich, D9891), 5000 IU/mL penicillin, 5 mg/mL streptomycin, 2 mM glutamine, and 10% (v/v) fetal bovine serum (Sigma), at 37 °C and 5% CO_2_.

FugeneHD (Promega) was used for both transient transfections and for integration. For integration in the HEK293T landing pad, 5x10^4^ cells seeded in 12-well dishes the day before were transfected with 0.1 μg of the pNLS-Bxb1-recombinase vector, mixed with 0.4 μg of the UBQLN2 plasmid, 40 μL OptiMEM (Gibco) and 1.6 μL FugeneHD. After two days, 10 nM of AP1903 (MedChemExpress) and 2 μg/mL doxycycline were added to select for recombinant cells for two days. The cells were used directly for western blotting, microscopy and flow cytometry after another two days of culturing in DMEM without AP1903 but supplemented with 2 µg/mL doxycycline.

The E6AP CRISPR/Cas9 KO plasmid (sc-401632, Santa Cruz) and E6AP HDR plasmid (sc-401632-HDR, Santa Cruz) were used to generate HEK293T landing pad E6AP KO cells, following the manufacturer’s instructions. A mixture of three sgRNAs were used. The sgRNA sequences were: GCTTACCTTGAGAACTCGAA, GACTTACTTAACAGAAGAGA, and GGGCACCTTTCGAGTTCTCA. Knockout was confirmed by western blotting.

### Denaturing immunoprecipitation

Denaturing immunoprecipitation was performed as described before ^24^. Briefly, at least 10^6^ HEK293T cells were seeded in 10-cm dishes. On the following day, the cells were transiently transfected with 4 µg of Strep-myc-ubiquitin and 4 µg of the different target constructs. One day after transfection, the cells were treated with bortezomib (LC Laboratories) at a concentration of 10 μM in serum-free DMEM for 16 hours. The cells were washed with PBS and harvested in 300 μL lysis buffer A (30 mM Tris/HCl pH 8.1, 2 mM EDTA, 100 mM NaCl and 0.2 mM PMSF). The samples were sonicated (on ice) three times for 10 seconds. Subsequently, 75 μL SDS 8% (w/v) was added, and the samples were boiled for 10 minutes. Then, 1125 μL lysis buffer A with 2.5% (v/v) Triton X-100 was added. After 30 minutes incubation on ice, the samples were centrifuged (16,000 g) for 60 min at 4 °C. Then, 20 μL of 50% slurry of anti-HA resin (Sigma) or myc-Trap beads (ChromoTek) in lysis buffer A was added to the cell extracts. After overnight tumbling at 4 °C, the beads were washed four times in 500 μL lysis buffer A containing 1 % (v/v) Triton X-100 and once with 500 μL lysis buffer A (without Triton X-100). Finally, 40 μL SDS sample buffer was added to the beads. The samples were boiled for 5 minutes and analyzed by electrophoresis.

### SDS-PAGE and blotting

Electrophoresis was performed using either 8 or 12.5% (w/v) acrylamide gels at 125-150 V. Semi-dry western blotting to 0.2 μm nitrocellulose membranes (Advantec) was performed using a constant current of 100 mAmp per gel for 60-90 min. The membranes were blocked with 5% (w/v) skimmed milk powder in PBS containing 0.1% Tween-20 and 5 mM NaN_3_ for at least 30 minutes at room temperature. The primary antibodies were: anti-HA (Roche, 1:3000, 11867423001), anti-myc (ChromoTek, 1:1000, AB_2631398), anti-E6AP (Santa Cruz, 1:500, sc-166689), anti-GFP (ChromoTek, 1:1000, 3H9), anti-RFP/mCherry (ChromoTek, 1:1000, 6G6). The secondary antibodies, all conjugated to HRP, were anti-mouse IgG (Dako, 1:5000, P0260) and anti-rat IgG (Thermo Fisher Scientific, 1:5000, 31470). Amersham ECL detection reagent (GE Healthcare) and a ChemiDoc Imaging System (BioRad) were used for development.

### Fluorescence microscopy

Live HEK293T cells expressing C-terminally tagged GFP-constructs, and mCherry from the landing pad integration site, were imaged directly using a Zeiss AxioVert.A1 microscope equipped with an AxioCam ICm 1 rev. camera. Excitation of EGFP and mCherry were at 475 nm and 590 nm, respectively. ImageJ v. 1.51j8 was used for processing the recorded images.

### Flow cytometry

For flow cytometry, dislodged cells were first washed in PBS by centrifugation, resuspended in 2 % (v/v) fetal bovine serum in PBS, and filtered through a 50 µm mesh filter. The cells were then applied to a BD FACSJazz (BD Biosciences) flow cytometer. Data collection and analyses were performed using FlowJo v.10.8.1 (BD), with the following gate settings: Live cells, singlet cells, BFP negative and mCherry positive. A minimum of 35,000 cells were analyzed. An example of the gating strategy is provided in the supplemental material (**Fig. S7**).

### Mass spectrometry data acquisition and analysis

HA tagged UBQLN2 variants were purified by denaturing immunoprecipitation using anti-HA resin (Sigma) as described above. Three separate biological repeats (independent transfections) per condition were analyzed in parallel. Purified proteins were fractionated on 4-12% polyacrylamide Bis-Tris NuPAGE gels (Thermofisher Scientific) using MOPS running buffer. After Coomassie staining and destaining, gel pieces were excised and tryptic peptides extracted as previously described ^70^. Peptide concentrations were normalized by UBQLN2 input and resuspended in 25-125 µL 0.1% TFA with 0.5% acetic acid. Mass spectrometry data acquisition was conducted by the MRC Protein Phosphorylation Unit proteomics facility at the University of Dundee. Briefly: 2 µL of each peptide sample derived from “Unmodified” UBQLN2 bands and 10 µL of the “Modified” forms, was analysed by LC-MS/MS using a Thermo Dionex Ultimate 3000 RSLC Nano liquid chromatography instrument coupled to an Orbitrap Exploris 240 (ThermoFisher Scientific, San Jose, CA) mass spectrometer operating in positive mode. HPLC liquid phase consisted of 0.1% formic acid as buffer A and 80% acetonitrile with 0.08% formic acid as buffer B and the solid phase comprised a C18 trap column in tandem with an EASY-Spray analytical column (C18, 2 μM, 75 μm x 50 cm) with an integrated nano electrospray emitter. Peptides were separated using a 70-minute method with a gradient from 3% to at least 95% of 80% acetonitrile with 0.08% formic acid at 45 °C. Eluted peptides were analysed with a data dependent method employing a 2 second cycle time; 2 kV spray voltage, 280 °C transfer tube temperature using a full precursor scan of 375-1500 at 60000 resolution and max injection time of 25 ms, followed by top N HCD MS/MS with minimum intensity of 5000, 1.2 m/z isolation window, charge state 2-6, NCE of 30, 100 ms max fill time, and 15000 resolution. A 30 second dynamic exclusion was applied. Further details of the LC-MS/MS settings can be obtained within the raw data files which have been deposited with the ProteomeXchange Consortium via the PRIDE ^71^ with the identifier PXD059465 (Username: Password: zPWvkEFSEK7s).

MS raw data files were processed by MaxQuant ^72^ version 2.1.3.0. All runs included variable modifications for GlyGly(K), LeuArgGlyGly(K), Phospho(ST), oxidised methionine and acetylated protein N-termini. Carbamidomethyl-C was fixed. FDR filtering was 1% at all levels. Other parameters were default with the exception of missed cleavages being set to 5, maximum peptide mass 9600 Da and minimum modified peptide Andromeda score of 30. Data were searched against the sequences of the exogenous UBQLN2 constructs that were duplicated with N-terminal fusion of ubiquitin in an attempt to identify N-terminal ubiquitylation. The entire UniProt Human proteome database (downloaded 22/05/2024) in addition to the in-built contaminants list were also searched. Evidence for lysine ubiquitylation was based on reported modified peptides containing the GlyGly(K) or the missed cleavage LeuArgGlyGly(K) adducts. See **SupplementalFile** for details.

## Acknowledgements

We thank Anne-Marie Lauridsen for technical assistance. We acknowledge the support from the FACS and computing core facilities at the Biotech Research & Innovation Centre and Department of Biology, University of Copenhagen. Fig. 3B and Fig. 5A were created with BioRender.com.

## Ethics approval and consent to participate

Not applicable.

## Consent for publication

Not applicable.

## Supplemental material

This article includes the following supplemental information.

- Supplemental figures (SupplementalFigures.pdf).
- Supplemental datasets (SupplementalFile.xlsx).

## Data availability

Mass spectrometry data files have been deposited with the ProteomeXchange Consortium (PXD059465, Username: Password: zPWvkEFSEK7s). All other data generated are included in the figures and supplemental files.

## Accession codes

- UBQLN2, Q9UHD9
- E6AP (UBE3A), Q05086

## Author contributions

M.G.-T., C.K., P.E., M.H.T., and M.A. performed the experiments. M.H.T., K.H., K.L.-L., W.B., and R.H.-P. analyzed the data. K.H., K.L.-L., W.B., and R.H.-P. conceived the study. M.G.-T., W.B., and R.H.-P. wrote the paper.

