## Supplemental figures for "The importance of UBQLN2 ubiquitylation for its turnover and localization"

*Supplemental material*

**Fig. S1**. *Sequence alignment of the human UBQLN paralogs.* p.2
**Fig. S2**. *UBQLN2 mass spectrometry data.* p.3
**Fig. S3**. *MS/MS spectra for selected UBQLN2 peptides.* p.5
**Fig. S4**. *MS/MS spectra for selected ubiquitin peptides.* p.6
**Fig. S5**. *The R → K and R → Q UBQLN2 variants.* p.7
**Fig. S6**. *Ubiquitin independent proteasomal degradation of UBQLN2 ΔUBL.* p.8
**Fig. S7**. *Gating strategy for flow cytometry.* p.9

```
                                                       ↓ ↓                     ↓
UBQLN1   -MAESGESGGPPGSQDSAAGAEGAGAPAAAASAEPKIMKVTVKTPKEKEEFAVPENSSVQQFKEEISKRFKSHTDQLVLI
UBQLN2   -MAENGESSGPPRPSRGPAAAQGS----AAAPAEPKIIKVTVKTPKEKEEFAVPENSSVQQFKEAISKRFKSQTDQLVLI
UBQLN4   MAEPSGAETRPP-------------------------IRVTVKTPKDKEEIVICDRASVKEFKEEISRRFKAQQDQLVLI
UBQLN3   -MAKGGEALPQGSP---------------APVQDPHLIKVTVKTPKDKEDFSVTDTCTIQQLKEEISQRFKAHPDQLVLI

UBQLN1   FAGKILKDQDTLSQHGIHDGLTVHLVIKTQNRPQDHSAQQTN----------------------------------TAGS
UBQLN2   FAGKILKDQDTLIQHGIHDGLTVHLVIKSQNRPQGQSTQPSN----------------------------------AAGT
UBQLN4   FAGKILKDGDTLNQHGIKDGLTVHLVIKTPQKAQDPAAATASSPSTPDPASAPSTTPASPATPAQPSTSGSASSDAGSGS
UBQLN3   FAGKILKDPDSLAQCGVRDGLTVHLVIKRQHRAMGNECPAAS----------------------------------VPTQ

UBQLN1   NVTTSSTPNSNSTSGSATSNPFGLGGLGGLAGLSSLGLNTTNFSELQSQMQRQLLSNPEMMVQIMENPFVQSMLSNPDLM
UBQLN2   NTTSASTPRSNSTPISTNSNPFGLGSLGGLAGLSSLGLSSTNFSELQSQMQQQLMASPEMMIQIMENPFVQSMLSNPDLM
UBQLN4   RRSSGGGPSPGAGEGSPSATASILSGFGGILGLGSLGLGSANFMELQQQMQRQLMSNPEMLSQIMENPLVQDMMSNPDLM
UBQLN3   GPSPGSLPQPSSIYPADGPPAFSLG---LLTGLSRLGLAYRGFPDQPSSLMRQHVSVPEFVTQLIDDPFIPGLLSNTGLV

UBQLN1   RQLIMANPQMQQLIQRNPEISHMLNNPDI RQTLELARNPAMMQEMMRNQDRALSNLESIPGGYNA RRMYTDIQEPMLS
UBQLN2   RQLIMANPQMQQLIQRNPEISHLLNNPDIMRQTLEIARNPAMMQEMMRNQDLALSNLESIPGGYNALRRMYTDIQEPMLN
UBQLN4   RHMIMANPQMQQLMERNPEISHMLNNPELMRQTMELARNPAMMQEMMRNQDRALSNLESIPGGYNALRRMYTDIQEPMFS
UBQLN3   RQLVLDNPHMQQLIQHNPEIGHILNNPEIMRQTLEFLRNPAMMQEMIRSQDRVLSNLESIPGGYNVLCTMYTDIMDPMLN

UBQLN1   AAQEQFGGNPFASLVSNTSSGEGSQPSRTENRDPLPNPWA----------------------------------------
UBQLN2   AAQEQFGGNPFASVGSSSSSGEGTQPSRTENRDPLPNPWA----------------------------------------
UBQLN4   AAREQFGNNPFSSLAGNSDSSS-SQPLRTENREPLPNPWS----------------------------------------
UBQLN3   AVQEQFGGNPFATATTDNATTTTSQPSRMENCDPLPNPWTSTHGGSGSRQGRQDGDQDAPDIRNRFPNFLGIIRLYDYLQ

UBQLN1   ----------------------------------PQTSQSSSASSG-----------TASTVGGTTGSTASGTSGQSTTA
UBQLN2   ----------------------------------PPPATQSSATTS-----------TTTSTGSGSGNSSSNATGNTVAA
UBQLN4   ----------------------------------PSPPTSQAPGSG-----------GEGTGGSGTSQVHPTVSNPFGIN
UBQLN3   QLHENPQSLGTYLQGTASALSQSQEPPPSVNRVPPSSPSSQEPGSGQPLPEESVAIKGRSSCPAFLRYPTENSTGQGGDQ

UBQLN1   PNLVPGVGASMFNTPGMQSLLQQITENPQLMQNMLSAPYMRSMMQSLSQNPDLAAQMMLNNPLFAGNPQLQEQMRQQLPT
UBQLN2   ANYVA----SIFSTPGMQSLLQQITENPQLIQNMLSAPYMRSMMQSLSQNPDLAAQMMLNSPLFTANPQLQEQMRPQLPA
UBQLN4   AASLG---SGMFNSPEMQALLQQISENPQLMQNVISAPYMRSMMQTLAQNPDFAAQMMVNVPLFAGNPQLQEQLRLQLPV
UBQLN3   DGAGKSSTGHSTNLPDLVSGLGDSANRVPFAPLSFSPTAAIPGIPEPPWLPSPAYPRSLRPDGMNPAPQLQDEIQPQLPL

UBQLN1   FLQQMQNPDTLSAMSNPRAMQALLQIQQGLQTLATEAPGLIPGFTPGLGALGS---------------------------
UBQLN2   FLQQMQNPDTLSAMSNPRAMQALMQIQQGLQTLATEAPGLIPSFTPGVGVGVLGTAIGPVGPVTPIGPIGPIVPFTPIGP
UBQLN4   FLQQMQNPESLSILTNPRAMQALLQIQQGLQTLQTEAPGLVPSLGSFGISRTPAPSAG----------------------
UBQLN3   LMHLQ------AAMANPRALQALRQIEQGLQVLATEAPRLLLWFMPCLAGTGSVAGGIE---------------------

UBQLN1   -----------------TGGSSGTNGSNATPSENTSPTAGTTEPGHQQFIQQMLQALAGVNP-QLQNPEVRFQQQLEQLS
UBQLN2   IGPIGPTGPAAPPGSTGSGGPTGPTVSSAAPSETTSPTS--ESGPNQQFIQQMVQALAGANAPQLPNPEVRFQQQLEQLN
UBQLN4   -----------------SNAGSTPEAPTSSPATPATSSPTGASSAQQQLMQQMIQLLAGSGNSQVQTPEVRFQQQLEQLN
UBQLN3   ----------------SREDPLMSEDPLPNPPPEVFPALDSAELGFLSPPFLHMLQDLVSTNPQQLQPEAHFQVQLEQLR

UBQLN1   AMGFLNREANLQALIATGGDINAAIERLLGSQPS   589
UBQLN2   AMGFLNREANLQALIATGGDINAAIERLLGSQPS   624
UBQLN4   SMGFINREANLQALIATGGDINAAIERLLGSQLS   601
UBQLN3   SMGFLNREANLQALIATGGDVDAAVEKLRQS---   655
```

**Fig. S1** – *Sequence alignment of the human UBQLN paralogs.*

Sequence alignment using ClustalW of human UBQLN1, UBQLN2, UBQLN3 and UBQLN4. Identical (black) and homologous (grey) residues have been shaded. All lysine (red), arginine (green) and cysteine (yellow) residues are colored. The arrows mark the lysine residues (K41, K43 and K66) that were found to be ubiquitylated in wild-type UBQLN2. The UBL domain is boxed. Note the reduced contents of lysine and cysteine residues in UBQLN1, UBQLN2, and UBQLN4.

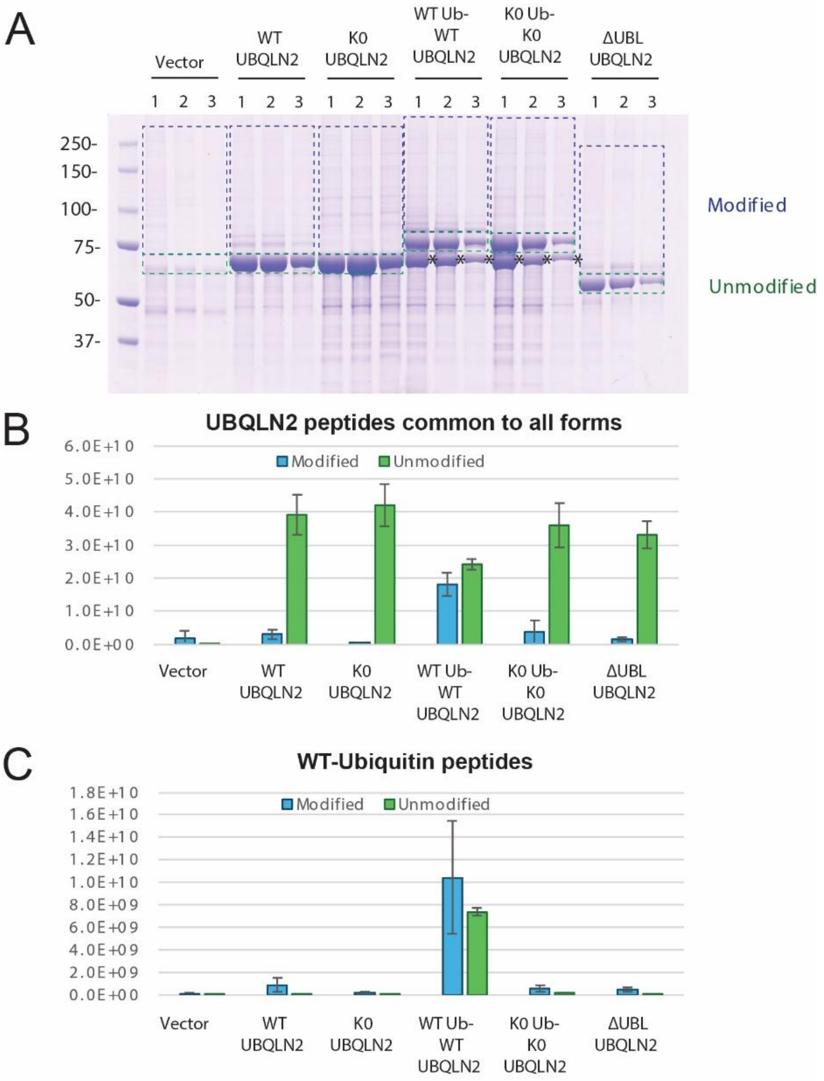


D

**Ubiquitin-modified peptides**

| | Vector | WT-UBQLN2 | K0-UBQLN2 | WT-Ub-WT-UBQLN2 | K0-Ub-K0-UBQLN2 | DeltaUBL-UBQLN2 |
|---|---|---|---|---|---|---|
| K6-Ubiquitin | 0.00 | 0.30 | 0.00 | 5.05 | 0.60 | 0.04 |
| K11 -Ubiquitin | 0.01 | 0.01 | 0.02 | 5.43 | 0.51 | 0.02 |
| K27 -Ubiquitin | 0.00 | 0.18 | 0.00 | 5.82 | 0.00 | 0.00 |
| K29 -Ubiquitin | 0.00 | 0.10 | 0.02 | 5.86 | 0.01 | 0.01 |
| K48 -Ubiquitin | 0.03 | 0.60 | 0.17 | 4.41 | 0.60 | 0.18 |
| K63 -Ubiquitin | 0.00 | 0.19 | 0.06 | 4.94 | 0.58 | 0.24 |
| K31 -UBQLN2 | 0.00 | 2.35 | 0.00 | 3.65 | 0.00 | 0.00 |
| K41 -UBQLN2 | 0.00 | 3.10 | 0.00 | 2.90 | 0.00 | 0.00 |
| K43 -UBQLN2 | 0.00 | 2.38 | 0.00 | 3.62 | 0.00 | 0.00 |
| K58 -UBQLN2 | 0.00 | 0.39 | 0.00 | 5.61 | 0.00 | 0.00 |
| K66 -UBQLN2 | 0.00 | 2.17 | 0.00 | 3.83 | 0.00 | 0.00 |
| K79 -UBQLN3 | 0.00 | 0.46 | 0.00 | 5.54 | 0.00 | 0.00 |

**Fig. S2** – *UBQLN2 mass spectrometry data.*

(A) Coomassie-stained SDS-PAGE gel fractionating immunoprecipitations of over-expressed HA-tagged UBQLN2 constructs or empty vector control. Samples were prepared in triplicate, and regions corresponding to unmodified and modified forms of UBQLN2 or N-terminally fused Ub-UBQLN2 were excised as shown. Asterisks (*) indicate the products of internal initiations of UBQLN2 from the N-terminally fused Ub-UBQLN2 constructs. Tryptic peptides were analysed by LC-MS/MS with loading normalized to represent approximately equal UBQLN2 for each lane. (BC) Total intensity of peptides common to all exogenous UBQLN2 forms (B) and WT ubiquitin which could be either exogenous or endogenous (C) from the samples shown in panel (A). Columns are averages and error bars are one standard deviation. (D) Heatmap of intensity of the indicated ubiquitination-specific peptides relative to the average intensity for the same peptide across all samples (modified slices only). MS signal intensity data for GG-K and LRGG-K peptides diagnostic of ubiquitination of the same lysine site were aggregated. Values are based on the average intensity of the three replicates per condition.

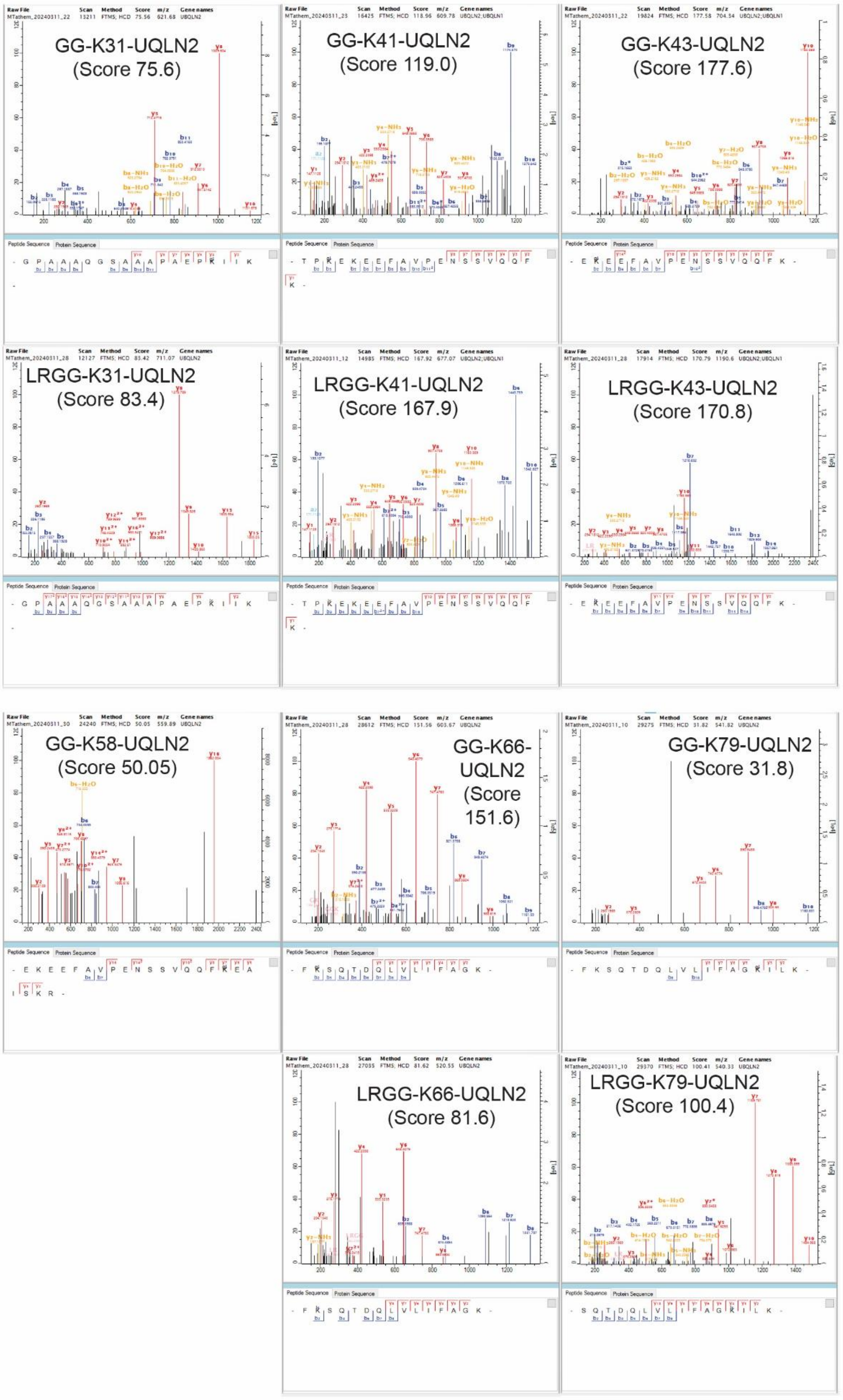


**Fig. S3** – *MS/MS spectra for the highest scoring peptides evidencing ubiquitination of UBQLN2*. Spectra for peptides carrying the GG and LRGG adducts for modification at the same site are shown where available.

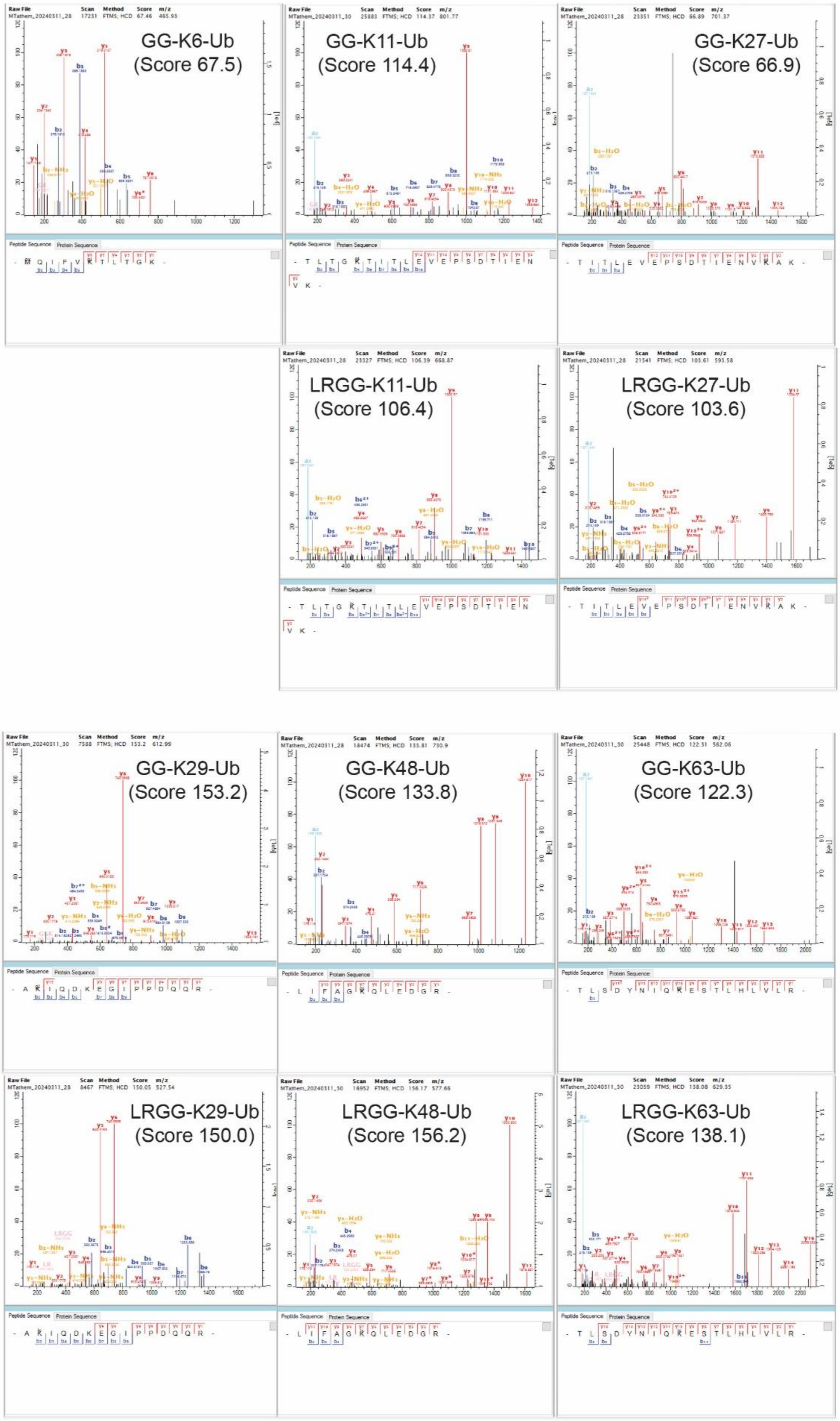


**Fig. S4** – *MS/MS spectra for the highest scoring peptides evidencing ubiquitination of ubiquitin itself.* Spectra for peptides carrying the GG and LRGG adducts for modification at the same site are shown where available.

```
UBQLN2 R>K   MAENGESSGPPRPSRGPAAAQGSAAAPAEPKIIKVTVKTPKEKEEFAVPENSSVQQFKEAISKRFKSQTDQLVLIFAGKI
UBQLN2 R>Q   MAENGESSGPPRPSRGPAAAQGSAAAPAEPKIIKVTVKTPKEKEEFAVPENSSVQQFKEAISKRFKSQTDQLVLIFAGKI
UBQLN2 WT    MAENGESSGPPRPSRGPAAAQGSAAAPAEPKIIKVTVKTPKEKEEFAVPENSSVQQFKEAISKRFKSQTDQLVLIFAGKI

UBQLN2 R>K   LKDQDTLIQHGIHDGLTVHLVIKSQNRPQGQSTQPSNAAGTNTTSASTPKSNSTPISTNSNPFGLGSLGGLAGLSSLGLS
UBQLN2 R>Q   LKDQDTLIQHGIHDGLTVHLVIKSQNRPQGQSTQPSNAAGTNTTSASTPQSNSTPISTNSNPFGLGSLGGLAGLSSLGLS
UBQLN2 WT    LKDQDTLIQHGIHDGLTVHLVIKSQNRPQGQSTQPSNAAGTNTTSASTPRSNSTPISTNSNPFGLGSLGGLAGLSSLGLS

UBQLN2 R>K   STNFSELQSQMQQQLMASPEMMIQIMENPFVQSMLSNPDLMKQLIMANPQMQQLIQKNPEISHLLNNPDIMKQTLEIAKN
UBQLN2 R>Q   STNFSELQSQMQQQLMASPEMMIQIMENPFVQSMLSNPDLMQQLIMANPQMQQLIQQNPEISHLLNNPDIMQQTLEIAQN
UBQLN2_WT    STNFSELQSQMQQQLMASPEMMIQIMENPFVQSMLSNPDLMRQLIMANPQMQQLIQRNPEISHLLNNPDIMRQTLEIARN

UBQLN2 R>K   PAMMQEMMKNQDLALSNLESIPGGYNALKKMYTDIQEPMLNAAQEQFGGNPFASVGSSSSSGEGTQPSKTENKDPLPNPW
UBQLN2 R>Q   PAMMQEMMQNQDLALSNLESIPGGYNALQQMYTDIQEPMLNAAQEQFGGNPFASVGSSSSSGEGTQPSQTENQDPLPNPW
UBQLN2 WT    PAMMQEMMRNQDLALSNLESIPGGYNALRRMYTDIQEPMLNAAQEQFGGNPFASVGSSSSSGEGTQPSRTENRDPLPNPW

UBQLN2 R>K   APPPATQSSATTSTTTSTGSGSGNSSSNATGNTVAAANYVASIFSTPGMQSLLQQITENPQLIQNMLSAPYMKSMMQSLS
UBQLN2 R>Q   APPPATQSSATTSTTTSTGSGSGNSSSNATGNTVAAANYVASIFSTPGMQSLLQQITENPQLIQNMLSAPYMQSMMQSLS
UBQLN2 WT    APPPATQSSATTSTTTSTGSGSGNSSSNATGNTVAAANYVASIFSTPGMQSLLQQITENPQLIQNMLSAPYMRSMMQSLS

UBQLN2 R>K   QNPDLAAQMMLNSPLFTANPQLQEQMKPQLPAFLQQMQNPDTLSAMSNPKAMQALMQIQQGLQTLATEAPGLIPSFTPGV
UBQLN2 R>Q   QNPDLAAQMMLNSPLFTANPQLQEQMQPQLPAFLQQMQNPDTLSAMSNPQAMQALMQIQQGLQTLATEAPGLIPSFTPGV
UBQLN2 WT    QNPDLAAQMMLNSPLFTANPQLQEQMRPQLPAFLQQMQNPDTLSAMSNPRAMQALMQIQQGLQTLATEAPGLIPSFTPGV

UBQLN2 R>K   GVGVLGTAIGPVGPVTPIGPIGPIVPFTPIGPIGPIGPTGPAAPPGSTGSGGPTGPTVSSAAPSETTSPTSESGPNQQFI
UBQLN2 R>Q   GVGVLGTAIGPVGPVTPIGPIGPIVPFTPIGPIGPIGPTGPAAPPGSTGSGGPTGPTVSSAAPSETTSPTSESGPNQQFI
UBQLN2 WT    GVGVLGTAIGPVGPVTPIGPIGPIVPFTPIGPIGPIGPTGPAAPPGSTGSGGPTGPTVSSAAPSETTSPTSESGPNQQFI

UBQLN2 R>K   QQMVQALAGANAPQLPNPEVRFQQQLEQLNAMGFLNREANLQALIATGGDINAAIERLLGSQPS   624
UBQLN2 R>Q   QQMVQALAGANAPQLPNPEVRFQQQLEQLNAMGFLNREANLQALIATGGDINAAIERLLGSQPS   624
UBQLN2 WT    QQMVQALAGANAPQLPNPEVRFQQQLEQLNAMGFLNREANLQALIATGGDINAAIERLLGSQPS   624
```

**Fig. S5** – *The R → K and R → Q UBQLN2 variants.*

Sequence alignment using ClustalW of wild-type, R → K and R → Q human UBQLN2 variants. All lysine residues are colored red.

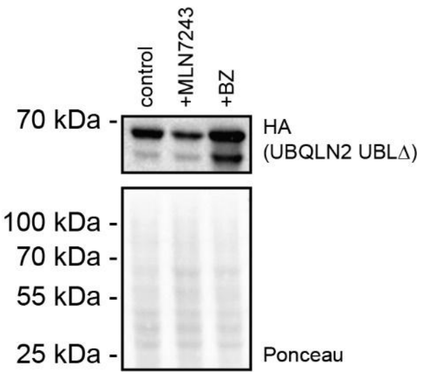


**Fig. S6** – *Ubiquitin independent proteasomal degradation of UBQLN2 ΔUBL.*
HEK293T cells transiently transfected to express the indicated UBQLN2-HA variant were treated with 10 μM of the proteasome inhibitor bortezomib (BZ) for 16 hours or 1 μM of the E1 inhibitor MLN7343 for 16 hours and analyzed by SDS-PAGE and western blotting. A Ponceau S staining is included as a control for equal loading.

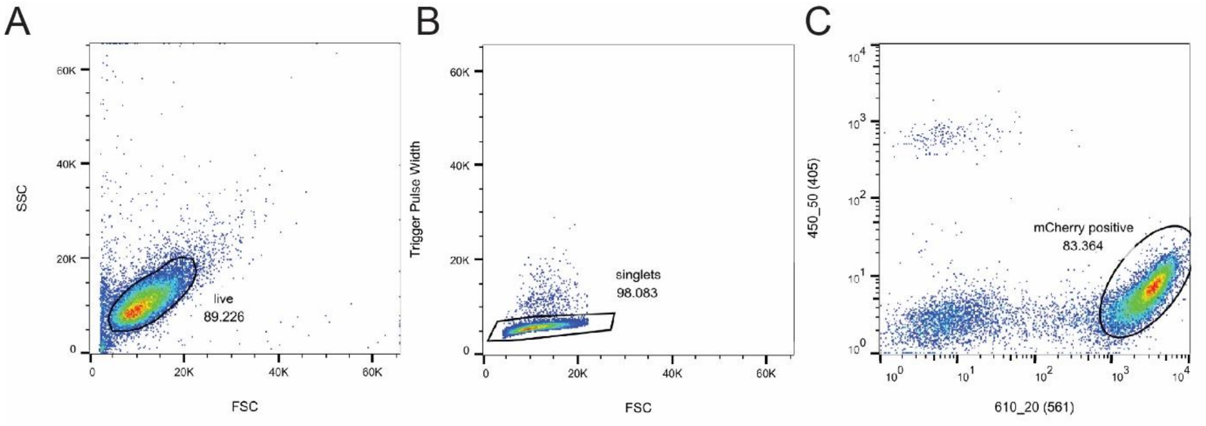


**Fig. S7** – *Gating strategy for flow cytometry.*

Example of the gating strategy used in flow cytometry. The cells were gated for: (A) live cells, based on the forward and side scattering, (B) singlets, based on the trigger pulse-width, and (C) mCherry positive and BFP negative signal, based on fluorescence.
